# A biointegrated living brain stimulator evokes specific neural signalling

**DOI:** 10.64898/2026.05.22.725279

**Authors:** Qilong Zhao, Qingbing Meng, Mingxing Peng, Zhonghua Lu, Zhiyuan Liu, Xiqun Jiang, Hairong Zheng, Xuemin Du

**Affiliations:** State Key Laboratory of Biomedical Imaging Science and System, Center for Intelligent Biomedical Materials and Devices, Shenzhen Institutes of Advanced Technology, Chinese Academy of Sciences, Shenzhen, China; Nano Science and Technology Institute, University of Science and Technology of China, Suzhou, China; University of Chinese Academy of Sciences, Beijing, China; State Key Laboratory of Biomedical Imaging Science and System, Guangdong Provincial Key Laboratory of Brain Connectome and Behavior, CAS Key Laboratory of Brain Connectome and Manipulation, The Brain Cognition and Brain Disease Institute, Shenzhen Technological Research Center for Primate Translational Medicine, Shenzhen Key Laboratory for Molecular Biology of Neural Development, Shenzhen-Hong Kong Institute of Brain Science, Shenzhen Institutes of Advanced Technology, Chinese Academy of Sciences, Shenzhen, China; CAS Key Laboratory of Human-Machine Intelligence-Synergy Systems, Shenzhen Institutes of Advanced Technology, Chinese Academy of Sciences, Shenzhen, China; MOE Key Laboratory of High Performance Polymer Materials and Technology and State Key Laboratory of Analytical Chemistry for Life Science, School of Chemistry, Nanjing University, Nanjing, China; State Key Laboratory of Biomedical Imaging Science and System, Research Center for Advanced Detection Materials and Medical Imaging Devices, Shenzhen Institutes of Advanced Technology, Chinese Academy of Sciences, Shenzhen, China

## Abstract

Implanted brain stimulators play a crucial role in treating various neurological disorders, including Parkinson’s disease (PD), Alzheimer’s disease, epilepsy, and depression. However, none of the existing implanted brain stimulators can realize specific neuromodulation due to fundamental disparities in signal transmission between electrical signal-induced neuronal responses and neurotransmitter-evoked neural signalling in natural neural circuits, leading to persistent challenges in biosafety and therapeutic effectiveness. Inspired by the dopaminergic neural circuit, we report a biointegrated living brain stimulator (BBS) that integrates ferroelectric bioelectronics, dopaminergic cells, and a gelatin hydrogel matrix, enabling dopamine neurotransmitter-evoked neural signalling. In contrast to conventional brain stimulators, the BBS are capable of programmed secretion of physiological-level dopamine, specifically activating nigral dopamine pathways and restoring motor function in a rodent PD model. By integrating the advantages of both bioelectronics and medicine, this lifelike BBS offers a great promise for next-generation bioelectronics, medicine, and brain-machine interfaces.

---

Implanted brain stimulators capable of modulating cellular activity in focal brain regions through electrical impulses have significantly advanced the treatment of neurological disorders, including Parkinson’s disease (PD)^1^, Alzheimer’s disease^2^, epilepsy^3^, and depression^4, 5^. Conventional brain stimulators typically consist of a surgically implanted system incorporating a metallic electrode for delivering electrical stimulation, a co-implanted battery for power supply, a controller for signal modulation, and intricate wiring to interconnect these components^6^. Although such brain stimulators have demonstrated clinical outcomes in alleviating motion symptoms in patients with severe disabilities, they still suffer from complicated implantation systems, pronounced foreign-body responses, and inevitable activation of neighboring non-target neurons^7, 8^. Alternatively, emerging brain stimulators have been developed based on inorganic photoelectric materials, inorganic piezoelectric materials, or inorganic magnetoelectric materials^9-15^, which are capable of converting external near-infrared (NIR) light, magnetic fields, or acoustic waves into electrical signals, therefore holding a promise to electrically modulate the neural activity in a wireless and battery-free manner^16-18^. However, despite these technological advances, current implanted brain stimulators are unable to achieve specific neuromodulation due to fundamental disparities in signal transmission between electrical signal-induced neuronal responses and neurotransmitter-evoked neural signalling in natural neural circuits^19^, leading to persistent challenges in biosafety and therapeutic efficacy^20^.

In natural neural circuits, neurons communicate through chemical neurotransmitters^21^. Dopamine, a key signaling molecule in the brain, regulates neuroplasticity and modulates essential functions such as locomotion, memory, and learning^22-24^. Inspired by dopamine-evoked neural signalling, we report a living bioelectronic medicine-a biointegrated living brain stimulator (BBS)-that harnesses the advantages of bioelectronics and medicine, enabling controllable dopamine provision to evoke specific neural signalling (Fig. 1a). Distinguished from conventional brain stimulators, this lifelike BBS integrates three critical components: (1) flexible ferroelectric bioelectronics (biocompatible polydopamine-blended poly(vinylidene fluoride-*co*-trifluoro ethylene) film with micropyramid array on the surface) imparts bioelectrical and biomechanical cues for cellular modulation; (2) rat pheochromocytoma (PC12) cells serve as the living components for sustained dopamine secretion; (3) a gelatin hydrogel layer preserves cell viability and promotes integration with biological tissues. Leveraging on their synergistic interactions, the BBS enables controllable and physiological-level dopamine release (∼7 ng·mL^-1^, equivalent to the dose in a normal rat’s striatum)^25^ under NIR irradiation.

**Figure 1.**
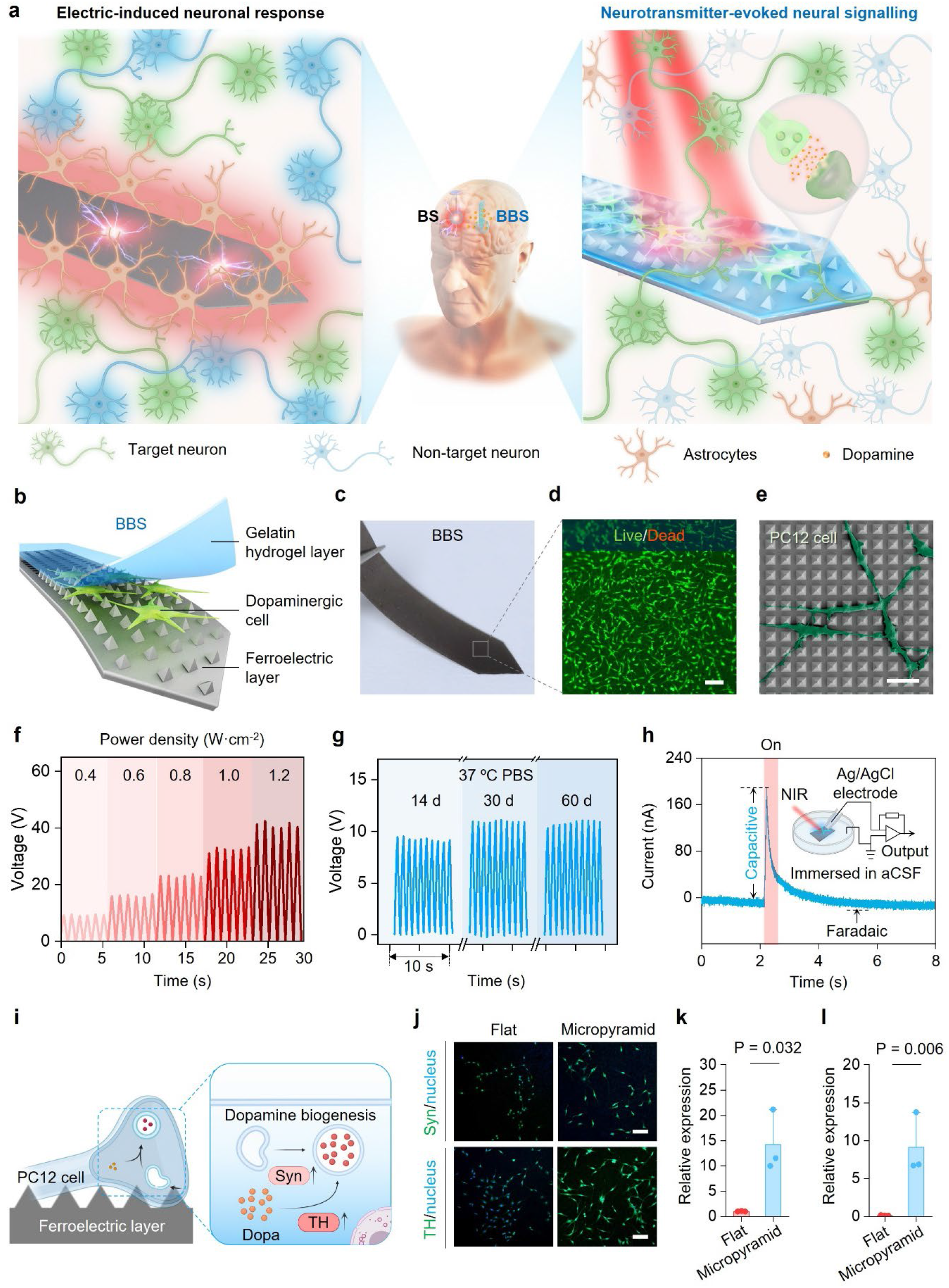
BBS Design. (**a**) Schematic illustrating fundamental disparities in signal transmission between electrical signal-induced neuronal responses for conventional brain stimulators (BS, left) and neurotransmitter-evoked neural signalling for the BBS (right). The BBS specifically activates nigral dopamine pathways, whereas the BS evokes non-specific cellular excitation in implantation sites. (**b**) Biointegrated configuration of the BBS. (**c**) A macroscopic photograph of the BBS, which is in the form of a flexible and uniform stripe with a sharp corner for facile insertion into brain. (**d**) Live/dead staining of the BBS after 1-day cell culture. PC12 cells show high viability (>98%) within the BBS. Scale bar: 100 μm. (**e**) Surface morphology of the BBS. PC12 cells (labelled in dark green) are well-stretched on the surface of the ferroelectric bioelectronics that possess ordered micro-pyramid array topography. Scale bar: 20 μm. (**f**) Representative curves showing the open-circuit voltages generated by the ferroelectric bioelectronics upon exposure to the NIR irradiation at different power densities (4, 6, 8, 10, or 12 mW·mm^-2^). (**g**) Stable open-circuit voltages generated by the ferroelectric bioelectronics in the aging test for 60 days under the same NIR irradiation (4 mW·mm^-2^, 1 Hz, 50% duty factor). (**h**) Representative curve showing the ionic current generated by the ferroelectric bioelectronics immersed in artificial cerebrospinal fluids (aCSF) under 808-nm NIR irradiation with a duration of 50 ms, measured by a patch-clamp setup. (**i**) Schematically illustrating the biomechanical modulation of the PC12 cells within the BBS. The BBS affords the biomechanical microenvironment for upregulating the expressions of TH and Syn of PC12 cells, thereby promoting the dopamine biogenesis. (**j**) Representative fluorescence images of the PC12 cells with stained Syn/TH and nucleus on the polydopamine-blended P(VDF-TrFE) films with smooth or micropyramid surfaces after 4-day culture. Scale bar: 100 μm. (**k, l**) Quantitative reverse transcription polymerase chain reaction (qRT-PCR) measurements of the Syn (**k**) and TH (**l**) mRNA expressions for the PC12 cells on the polydopamine-blended P(VDF-TrFE) films with smooth or micropyramid surfaces after 4-day culture. Syn: smooth vs micropyramid: P = 0.032 (*), TH: Syn: smooth vs micropyramid: P = 0.006 (**); n = 3 separate experiments. Statistical analyses into the data shown in **k** and **l** were performed using one-way analysis of variance (ANOVA) with two-tailed Student’s *t*-test.

This lifelike BBS achieves specific activation of dopaminergic signalling both in vitro and in vivo, a capability not attainable with existing brain stimulators. In a rodent PD model, the BBS not only restores motor function but also demonstrates no detectable side effects and negligible foreign-body responses even after long-term implantation (∼300 days). As a new class of therapeutic modality, such living bioelectronic medicine holds great promise for advancing next-generation brain-machine interfaces, intelligent biomedical devices, and biosafe, effective treatments for neurological disorders^26-30^.

## BBS Design

To construct the BBS, we sequentially integrate the ferroelectric bioelectronics, dopaminergic cells, and the gelatin hydrogel matrix (Fig. 1b). The ferroelectric bioelectronics are fabricated through a multi-step process, including the synthesis of polydopamine particles, dispersing the polydopamine particles into a P(VDF-TrFE) solution, casting the former precursor into a pre-made silicon mold with inverse micropyramid array, and finally poling the film (Fig. S1a). Polydopamine particles are selected as photothermal agents due to their superior biocompatibility, efficient light-to-thermal conversion under NIR irradiation, and excellent compatibility with P(VDF-TrFE)^31, 32^. P(VDF-TrFE) is incorporated for its unique ferroelectric properties and robust stability, enabling electric field generation^33-36^. Harnessing their combined advantages, the ferroelectric bioelectronics exhibit scalable fabrication, tissue-like flexibility, excellent biocompatibility, and tunable bioelectrical properties (Fig. 1c, d and Fig. S1b, c). The resulting cellular-sized micropyramid arrays on the ferroelectric bioelectronics surface impart biomechanical cues for subsequent cellular modulation (Fig. 1e and Fig. S1d). Second, the PC12 cells, as a typical class of dopaminergic cells, are seeded onto the surface of the ferroelectric bioelectronics to serve as the living components, capable of exocytosis of dopamine-containing synaptic vesicles essential for neurotransmitter-mediated signalling in dopaminergic neural circuit^23^. Third, gelatin hydrogel fibers with naturally derived composition and excellent biocompatibility^37^, are prepared and covered onto the surface of the cell-laden ferroelectric bioelectronics to preserve PC12 cell viability and facilitate seamless integration with biological tissues.

Upon exposure to NIR irradiation, the ferroelectric film within the BBS exhibits a rapid temperature rise, thus generating a volt-level electric field (Fig. S2). The magnitude of the NIR-induced electric field can be facilely tuned by adjusting polydopamine content, NIR power density, or irradiation frequency (Fig. 1f and Fig. S2b-h). Moreover, this capability remains stable, with no detectable decay after immersion in 37 ºC phosphate-buffered saline for 60 days (Fig. 1g). Notably, this ferroelectric film can generate NIR-induced electric fields even in artificial cerebrospinal fluids. Patch-clamp measurement demonstrates a transient peak current corresponding to capacitive effects, while Faradaic effects are negligible (Fig. 1h). The nanoampere-level capacitive current is proven sufficient and safe for electrical cell stimulation^15^. In addition, the BBS provides a favorable biomechanical microenvironment that promotes cell attachment and spreading, as evidenced by well-stretched PC12 cell morphologies on its surface (Fig. 1d and Fig. S1e). This biomechanical cue further activates biophysical signalling pathways in PC12 cells^38^, enhancing neuronal differentiation and upregulating neuroplasticity-related proteins, including tyrosine hydroxylase (TH) and synaptophysin (Syn), which are key factors involved in dopamine synthesis and synaptic vesicle formation, respectively^39^, compared to flat controls (Fig. 1i-l, Fig. S3a, and Fig. S4). By integrating superior bioelectrical and biomechanical cues with living cells, such lifelike features would impart a robust capability for BBS to modulate electrophysiological responses and exocytosis of PC12 cells.

### Programmable secretion of dopamine neurotransmitters

Electrophysiological modulation of dopaminergic cells can regulate the trafficking of dopamine-containing synaptic vesicles, a process critical for dopamine secretion (Fig. S3b) ^40^. To assess this capability, we evaluate the electrophysiological responses of PC12 cells within the BBS (Fig. 2a). Upon pulsed NIR irradiation, a substantial proportion of cells exhibits significantly increased intracellular Ca^2+^ fluorescence (Fig. 2b, c and Video 1). In contrast, PC12 cells on a control film lacking bioelectrical behavior do not respond to the same NIR irradiation, indicating that the NIR-induced bioelectrical, rather than photothermal effects, induces electrophysiological responses (Fig. 2b, d, Fig. S5, and Video 2). Notably, reliable cellular excitation is maintained even at lower power densities or higher frequencies, with minimal variation in the proportion of excitable cells (Fig. 2e, Fig. S6 and Videos 3-5). Importantly, PC12 cells within the BBS generate negligible levels of intracellular reactive oxygen species, which is ∼5 times lower than that on a photovoltaic Si film under similar NIR-induced excitation, thus preserving high viability (> 95%) even after 7 days of continuous NIR irradiation (Fig. S7, Fig. S8, and Videos 6-8). Whole-cell patch-clamp recordings further confirm that NIR irradiation elicits synchronous and significant depolarization of the cell membrane potential in the BBS (Fig. 2f-h and Fig. S9), whereas control samples show negligible changes (Fig. 2f). Moreover, repeated on/off cycles of NIR irradiation can induce reproducible excitation of PC12 cells (Fig. 2f), implying the programmable dopamine secretion capability of the BBS.

**Figure 2.**
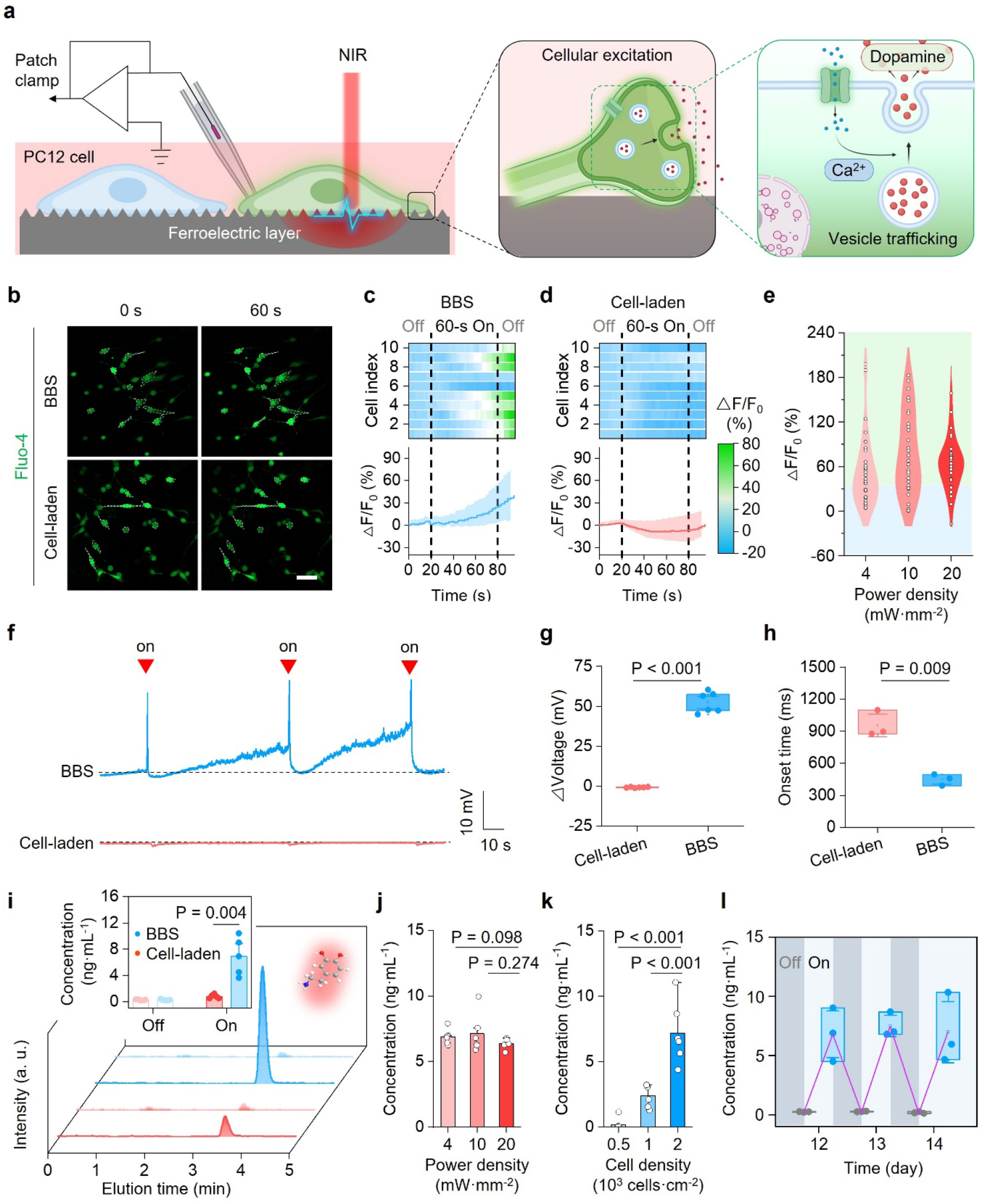
Programmable secretion of dopamine neurotransmitters. (**a**) Schematically illustrating the electrophysiological modulation of the PC12 cells within the BBS for programmable DA secretion. The BBS affords the bioelectrical cues for evoking the excitation of PC12 cells as exposure to NIR irradiation, therefore triggering the trafficking and exocytosis of the dopamine-containing synaptic vesicles. (**b**) Representative time-lapse fluorescence images of the PC12 cells stained with fluo-4 AM (an intracellular Ca^2+^ indicator) under NIR irradiation (power density: 20 mW·mm^-2^, frequency: 1 Hz, duty factor: 50%, duration: 1 min) within the BBS or cultured on the polydopamine-blended poly(vinyl chloride) films with micropyramid array topography yet without ferroelectric property (Cell-laden). Scale bar: 50 μm. (**c, d**) Normalized fluorescence changes (ΔF/F_0_) over time of each PC12 cell stained with the fluo-4 AM under the NIR irradiation within the BBS (**c**) or cultured on the cell-laden films (**d**). The proportion of excitatory cells (ΔF/F_0_ > 40%) within the BBS under the NIR irradiation is above 80%, while all the cells cultured on the cell-laden films show no change of fluorescence signals (ΔF/F_0_ > 20%). (**e**) Normalized fluorescence changes (ΔF/F_0_) for the PC12 cells within the BBS stained with the fluo-4 AM under NIR irradiation with varying power densities of 4, 10, or 20 mW·mm^-2^. n = 50 cells. (**f**) Representative curves showing the changes of cellular membrane potential of the PC12 cells within the BBS or cultured on the cell-laden films under the same NIR irradiation (power density: 4 mW·mm^-2^, duration: 500 ms). (**g**) Statistics of the changes of the membrane potential for the PC12 cells within the BBS or the cell-laden devices under NIR irradiation, showing an average amplitude of 51 ± 3 mV for the BBS group. BBS vs Cell-laden: P < 0.001 (***); n = 6 biologically independent experiments. (**h**) Statistics of the onset time for the PC12 cells to change their membrane potential within the BBS or the cell-laden devices under NIR irradiation, showing an average onset time of 449 ms for the BBS group. BBS vs Cell-laden: P = 0.009 (**); n = 3 spikes. (**i**) HPLC/MS chromatographs showing the contents of dopamine secreted by the BBS or the Control before or after NIR irradiation. The inset shows the statistics of dopamine concentrations released by the BBS or the cell-laden devices before and after NIR irradiation. NIR off: BBS vs Cell-laden: P = 0.486 (n. s.); NIR on: BBS vs Cell-laden: P = 0.004 (**). n = 5 separate experiments. (**j**) Dopamine secretion by the BBS with varying cell densities (500, 1000, or 2000 cells·cm^-2^) under NIR irradiation with a power density of 20 mW·mm^-2^, 1 Hz frequency, and 50% duty factor. n = 6 separate experiments. 500 cells·cm^-2^ vs 1000 cells·cm^-2^: P < 0.001 (***); 1000 cells·cm^-2^ 2000 cells·cm^-2^: P < 0.001 (***). (**k**) Dopamine secretion by the BBS with a cell density of 2000 cells·cm^-2^ under NIR irradiation with varying power densities (4, 10, or 20 mW·mm^-2^). 4 mW·mm^-2^ vs 20 mW·mm^-2^: P = 0.098 (n. s.); 10 mW·mm^-2^ vs 20 mW·mm^-2^, P = 0.274 (n. s.). n = 6 separate experiments. (**l**) Dopamine secretion by the BBS after culturing different days (12, 13, or 14 days). n = 3 separate experiments. Statistical analyses into the data shown in **g, h, i, j** and **k** were performed using one-way ANOVA with two-tailed Student’s *t*-test.

We next investigate the NIR-mediated programmable dopamine secretion from the BBS. In the absence of NIR stimulation, the BBS releases minimal dopamine (< 0.5 ng·mL^-1^) under resting status, as determined by high-performance liquid chromatography-tandem mass spectrometry (HPLC/MS, Fig. 2i, Fig. S10). Upon NIR irradiation, the BBS secrets dopamine at a physiological level (∼7 ng·mL^-1^), comparable to concentrations found in normal rat striatum, and markedly higher than those from a control cell-laden device (∼7-fold increase) or directly transplanted dopaminergic cells^25^ (∼2-fold increase, Fig. 2i). Dopamine release from the BBS remains consistent across varying NIR power densities, which agrees well with former cellular electrophysiological responses (Fig. 2j). Furthermore, by varying the density of PC12 cell, dopamine secretion can be programmed from ∼1 to ∼7 ng·mL^-1^ (Fig. 2k). Notably, the BBS maintains stable dopamine release at ∼7 ng·mL^-1^ from day 12 to 14, extending the release period to ∼9 times longer than existing dopamine delivery systems (∼36 hours, Fig. 2l)^41, 42^. This sustained, on-demand release avoids adverse effects associated with oxidation of pre-loaded dopamine to toxic dopaquinone and the risks of overdose, common in conventional dopamine-supplementation strategies^43^. Collectively, these results demonstrate that the BBS enables unprecedented, programmable provision of dopamine, supplying essential chemical neurotransmitters for specific neuromodulation and restoration of dopaminergic signalling in dopamine-deficit diseases.

### Specifically evoking dopaminergic signalling

We next examine whether the BBS can evoke dopaminergic signalling in vitro. The BBS is cocultured with primary rat-derived hippocampal neurons, which possess abundant dopamine receptors and are integral to dopaminergic circuit function (Fig. 3a)^44^. Hippocampal neurons extend neurites along the micropyramid array structures, forming dense neural networks facilitated by the unique biomechanical cues of the BBS (Fig. 3b, c and Fig. S11). This feature further upregulates the neuroplasticity-related proteins (i.e., Syn and TH), and promotes extensive contacts between PC12 cells and target neuronal neurites (Fig. 3d, e, Fig. S11c, and Fig. S12), both essential for neural signalling^45, 46^. To further assess dopamine receptor activation, hippocampal neurons are labeled with genetically encoded green-fluorescent G-protein-coupled receptor-activation-based dopamine (GRAB_DA_) sensors (Fig. S13). Upon NIR irradiation of the BBS, a significant increase in fluorescent intensity is observed, indicating dopamine receptor activation (Fig. 3f and Video 9). In contrast, control samples possessing solely biomechanical or biomechanical and bioelectrical cues do not elicit changes in the fluorescent intensity under identical NIR conditions (Fig. 3g, Fig. S13, and Videos 10, 11). These results indicate that the BBS enables specific activation of dopamine receptors in vitro, a hallmark of dopaminergic signalling^47^. Correspondingly, cocultured neurons exhibit robust increases in intracellular Ca^2+^ fluorescence under NIR irradiation, indicating neuronal excitation, whereas the control shows no such alterations (Fig. 3h, i, Fig. S14, and Videos 12-14).

**Figure 3.**
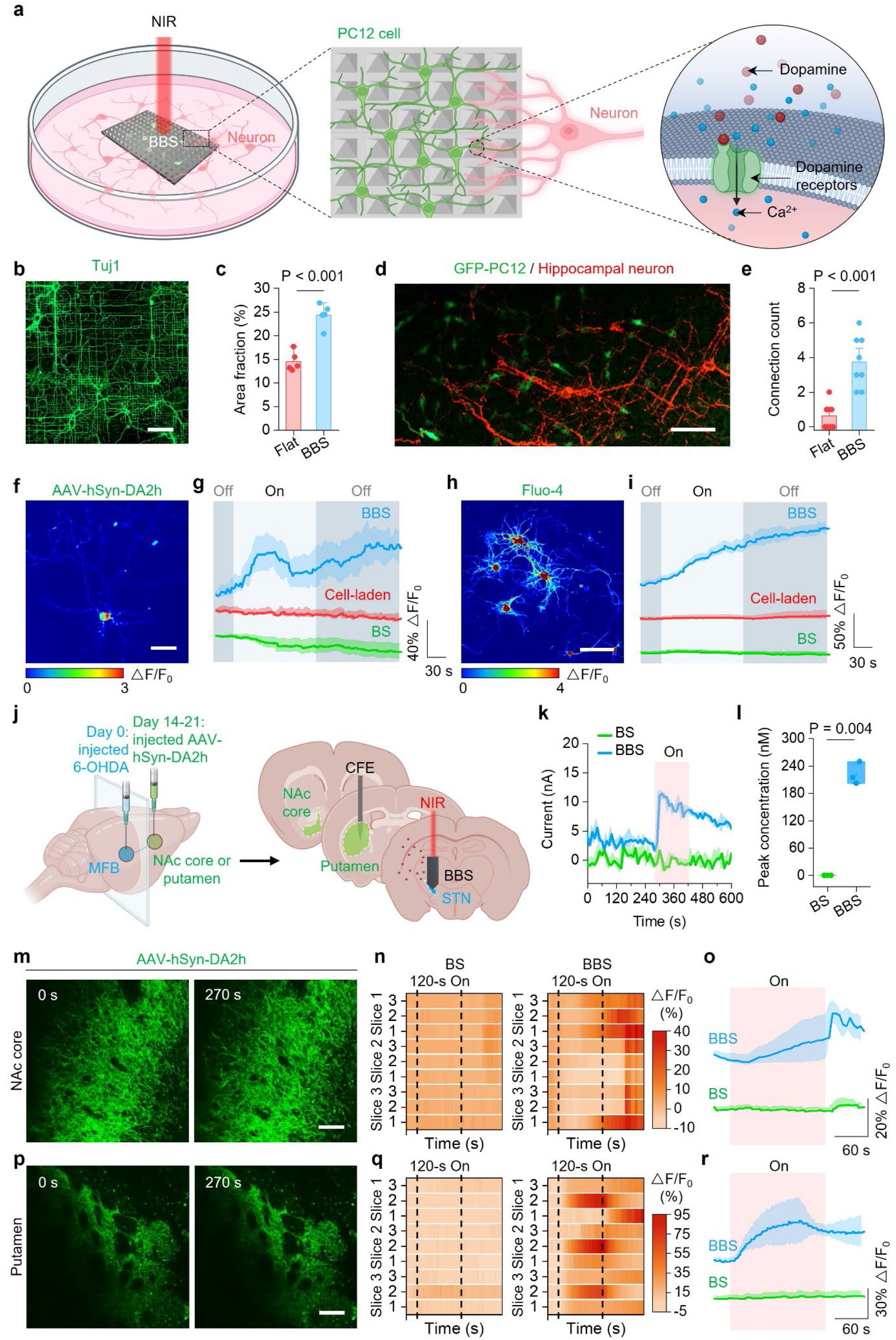
Specifically evoking dopaminergic signalling. (**a**) Schematically illustrating the interfacing and communication between the BBS and target neurons. The BBS forms a seamless interface with target neurons by forming vast contacts between the PC12 cells and the neurites of target neurons. Upon exposure to NIR irradiation, the BBS can specifically activate the post-synaptic dopamine and then elicit the excitation of target neurons through the programmable dopamine provision. (**b**) Representative fluorescence images of mouse-derived primary hippocampus neurons with stained Tuj1 on the surface of the BBS after 14-day culture. Scale bar: 100 μm. (**c**) Statistics of the area fraction (%) for the Tuj1-positive cells within the BBS or cultured on the flat ferroelectric films (Flat) after 14-day culture. Flat vs BBS: P < 0.001 (***); n = 5 separate experiments. (**d**) Representative fluorescence images showing the interface between the Tuj1-stained neurons (red) and the GFP-expressed PC12 cells (GFP-PC12, green) within the BBS after 14-day coculture. Scale bar: 100 μm. (**e**) Statistics of the intercellular connection density between PC12 cells and neurons within the BBS or the Flat after 14-day coculture. F-BBS vs BBS: P = 0.002 (**); n = 5 separate experiments. (**f**) Representative images of GRAB_DA_ sensor response to dopamine released by the BBS under NIR irradiation (power density: 20 mW·mm^-2^, frequency: 1 Hz, duty factor: 50%, duration: 2 min). Scale bar: 50 μm. (**g**) Average traces of normalized fluorescence changes (ΔF/F_0_) over time for the neurons expressing the GRAB_DA_ sensors co-incubated with the BBS, the cell-laden device formulated by seeding PC12 cells on the polydopamine-blended PVC films (Cell-laden), or the cell-free brain stimulator (ferroelectric layer alone, BS) under the same NIR irradiation (power density: 20 mW·mm^-2^, frequency: 1 Hz, duty factor: 50%, duration: 2 min). n = 10 regions. (**h**) Representative images of intracellular Ca^2+^ probe fluo-4 AM response to dopamine released by the BBS under NIR irradiation (power density: 20 mW·mm^-2^, frequency: 1 Hz, duty factor: 50%, duration: 2 min). Scale bar: 50 μm. (**i**) Average traces of normalized fluorescence changes (ΔF/F_0_) over time for the neurons stained with the fluo-4 AM co-incubated with the BBS, the cell-laden device (Cell-laden), or the cell-free brain stimulator (BS) under the same NIR irradiation (power density: 20 mW·mm^-2^, frequency: 1 Hz, duty factor: 50%, duration: 2 min). The fluorescence signal shows a significant rise for the BBS (ΔF/F_0_ > 200%), while the other two groups exhibit steady fluorescence signals (ΔF/F_0_ < 20%). n = 5 cells. (**j**) Experimental scheme of monitoring dopaminergic signalling. Dopamine delivered to target regions of dopamine signalling is monitored via square wave voltammetry (SWV) by using a functionalized carbon fiber electrode. (**k**) Average characteristic SWV peak current to dopamine over time for the BBS and cell-free brain stimulators (BS) before and after NIR irradiation. n = 3 separate experiments. (**l**) Statistic of peak dopamine concentrations in striatum delivered by the BBS or cell-free brain stimulator (BS) under NIR irradiation, which is determined by converting SWV peak current to dopamine concentration according to the calibration in vitro. BBS vs BS: P = 0.004 (**). n = 3 separate experiments. (**m**) Time-lapse fluorescence images for the neurons expressing the GRAB_DA_ sensors in the NAc core at acute brain slices from adult mice stimulated by the BBS or the cell-free control under NIR irradiation (power density: 3 mW·mm^-2^, frequency: 1 Hz, duty factor: 50%, duration: 2 min). (**n, o**) Normalized fluorescence changes (ΔF/F_0_) over time for each region (**n**) and average traces (**o**) of the neurons expressing the GRAB_DA_ sensors in the NAc core stimulated by the BBS or the cell-free brain stimulator (BS) under the same NIR irradiation. n = 9 regions from 3 mice. (**p**) Time-lapse fluorescence images for the neurons expressing the GRAB_DA_ sensors in the putamen at acute brain slices from adult mice stimulated by the BBS or the cell-free brain stimulator (BS) under NIR irradiation (power density: 3 mW·mm^-2^, frequency: 1 Hz, duty factor: 50%, duration: 2 min). (**q, r**) Normalized fluorescence changes (ΔF/F_0_) over time for each region (**q**) and average traces (**r**) of the neurons expressing the GRAB_DA_ sensors in the putamen stimulated by the BBS or the cell-free brain stimulator (BS) under the same NIR irradiation. n = 9 regions from 3 rats. Statistical analyses into the data shown in **c, e** and **l** were performed using one-way ANOVA with two-tailed Student’s *t*-test.

Consistent with in vitro results, the BBS also enables programmable dopamine secretion and activation of nigral dopaminergic signalling in vivo. To assess *in vivo* feasibility, we first quantify dopamine release from the BBS under NIR irradiation, accounting for attenuation by mouse’s skull tissues (Fig. S15a). Despite partial attenuation, the BBS maintains dopamine secretion at ∼6.2 ng mL^-1^ under NIR irradiation (Fig. S15b, c). This NIR-mediated dopamine release is further validated using a functionalized carbon fiber electrode implanted in the rat striatum, with the BBS positioned in the medial forebrain bundle (MFB, Fig. 3j and Fig. S15d-l). In brain slices from BBS-treated rats, dopamine signals increase sharply within ∼10 s of NIR exposure, whereas cell-free controls show no change (Fig. 3k). The peak dopamine concentration delivered to the target striatum reaches ∼220 nM (Fig. 3l), comparable to physiological levels observed during striatum activation (∼260 nM)^48^. To confirm specific activation of post-synaptic neurons, we monitor GRAB_DA_ sensor signals in the nucleus accumbens (NAc) and putamen, key targets of dopaminergic signalling^23^. NIR irradiation of the BBS elicits robust activation of dopamine receptors in these regions, while cell-free brain stimulators produce no detectable changes (Fig. 3m-r, Fig. S16, and Videos 15-18). By mimicking interneural communication within the natural dopaminergic neural circuit^23^, the BBS avoids off-target activation of neighboring neurons and enables reliable neuromodulation even when spatially separated from target neurons, a robust capability not achievable with conventional brain stimulators, which suffer from sharp electric signal attenuation at electrolyte-/cell-electrode interfaces^15^. These results highlight the potential of the BBS as a novel strategy for neuromodulation therapies through restoration of dopaminergic signalling.

### Reversing parkinsonian symptoms in rodent model

We next evaluate the therapeutic potential of the BBS for PD, one of the most predominant neurological disorders caused by defective dopaminergic signalling and still lacking effective approaches for precise modulation^49-52^. Briefly, a PD mouse model is established by bilateral injection of 6-hydroxydopamine (6-OHDA) into the MFB, resulting in complete ablation of dopaminergic neurons (Fig. 4a and Fig. S17a, b). The BBS is then implanted into the unilateral subthalamic nucleus of PD mice exhibiting characteristic symptoms, with precise placement confirmed by magnetic resonance imaging (MRI), which reveals negligible artifact (Fig. 4b, Fig. S17c, d). Beginning 14 days post-implantation, the BBS is irradiated with 808-nm NIR light for 3 consecutive days, demonstrating effective transcranial activation and biosafety for brain tissues (Fig. S17e-i)^53^. Notably, PC12 cells within the BBS can also maintain a high density (∼3×10^5^ cells·cm^-2^) after 21 days in vivo (Fig. S17j).

**Figure 4.**
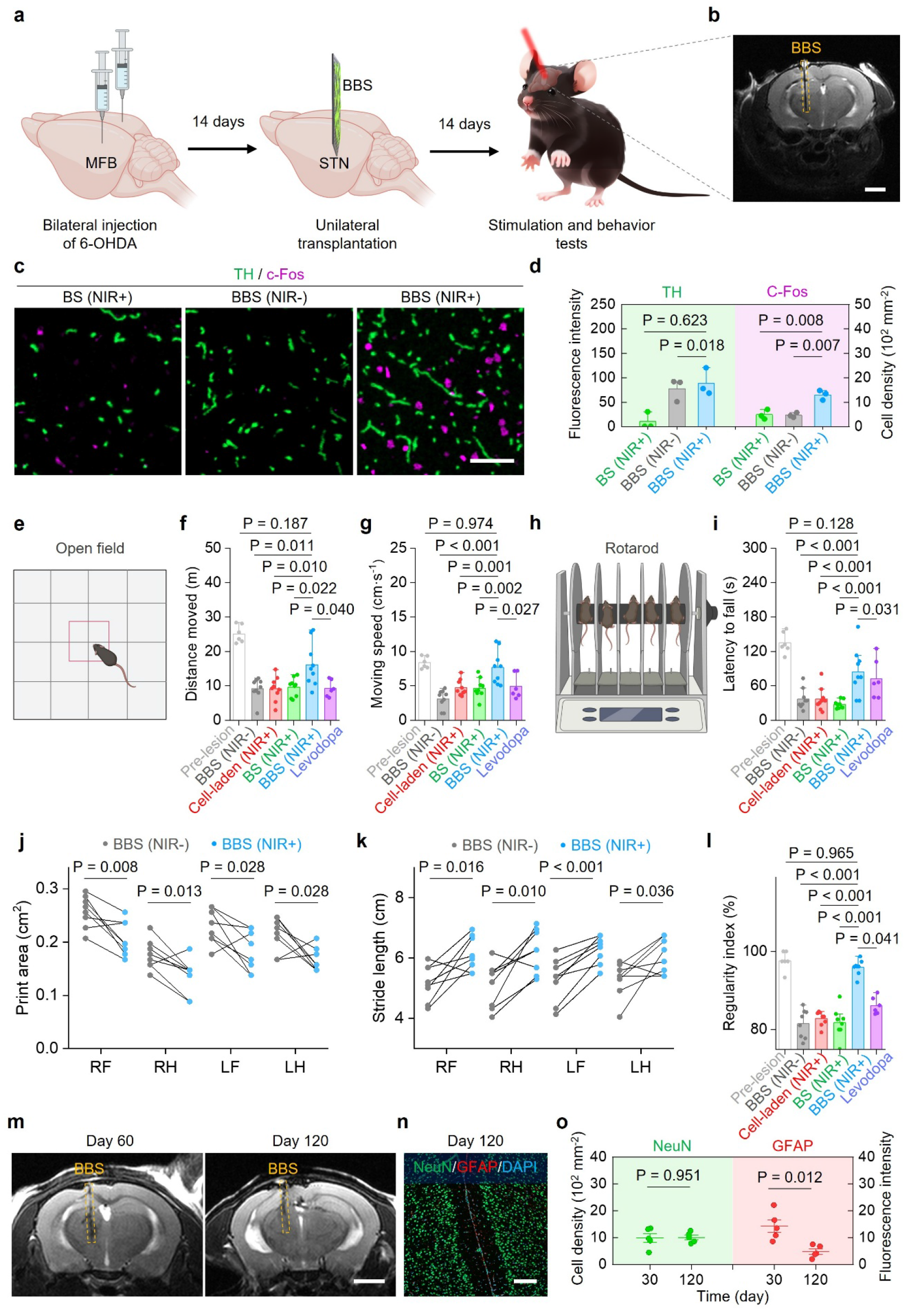
Reversing parkinsonian symptoms in rodent model. (**a**) Experimental scheme for BBS implantation and in vivo evaluation. (**b**) Magnetic resonance imaging (MRI) shows the fully implanted BBS (highlighted in a dashed square) in a mouse’s brain. Scale bar: 2 mm. (**c**) Representative fluorescence images of TH-positive and c-Fos-positive cells in the substantia nigra pars compacta contralateral and ipsilateral to the implantation. Scale bar: 50 μm. (**d**) Statistics of fluorescence intensity of TH-positive cells and cell densities of c-Fos-positive cells among different groups (BBS (NIR-): implanting the BBS yet absent of NIR irradiation, BS (NIR+): implanting the ferroelectric layer alone absent of PC12 cells and irradiated with NIR light, and BBS (NIR+): implanting the BBS and irradiated with NIR light). TH fluorescence intensity: BBS (NIR+) vs BBS (NIR-): P = 0.623 (n. s.), BBS (NIR+) vs BS (NIR+): P = 0.018 (*); c-Fos-positive cell density: BBS (NIR+) vs BBS (NIR-): P = 0.008 (**), BBS (NIR+) vs BS (NIR+): P = 0.007 (**). n = 3 separate experiments. (**e**) Experimental scheme of open field tests. (**f, g**) Statistics of total distance moved (**f**) and average moving speed (**g**) in the open field tests for the health mice (Pre-lesion), the PD mice implanted with the cell-laden devices (Cell-laden), the cell-free devices (BS), or the BBS before (NIR-) and after (NIR+) the NIR irradiation (frequency: 1 Hz, duty factor: 50%, power density: 20 mW·mm^-2^; 5-s on and 15-s off alternately performed for 24 recycles every day to individual subject), and the PD mice treated with levodopa. Total distance moved: BBS (NIR+) vs BBS (NIR-): P = 0.011 (*), BBS (NIR+) vs BS (NIR+): P = 0.022 (*), n = 9 mice; BBS (NIR+) vs Cell-laden (NIR+): P = 0.010 (*), n = 9 mice; BBS (NIR+) vs levodopa: P = 0.040 (*), 9 mice in the BBS (NIR+) group and 6 mice in the levodopa group; BBS (NIR+) vs Pre-lesion: P = 0.187 (n. s.), 9 mice in the BBS (NIR+) group and 6 mice in the pre-lesion group. Average moving speed: BBS (NIR+) vs BBS (NIR-): P < 0.001 (***), BBS (NIR+) vs BS (NIR+): P = 0.001 (**), n = 9 mice; BBS (NIR+) vs Cell-laden (NIR+): P = 0.002 (**), n = 9 mice; BBS (NIR+) vs levodopa: P = 0.027 (*), 9 mice in the BBS group and 6 mice in the levodopa group; BBS (NIR+) vs Pre-lesion: P = 0.974 (n. s.), 9 mice in the BBS (NIR+) group and 6 mice in the pre-lesion group. (**h**) Experimental scheme of rotarod tests. (**i**) Statistics of latency to fall in the rotarod for the health mice (Prelesion), the PD mice implanted with the cell-laden devices (Cell-laden), the cell-free devices (BS), or the BBS before (NIR-) and after (NIR+) the NIR irradiation, and the PD mice treated with levodopa. BBS (NIR+) vs BBS (NIR-): P < 0.001 (***), n = 9 mice; BBS (NIR+) vs BS (NIR+): P < 0.001 (***), n = 9 mice; BBS (NIR+) vs Cell-laden (NIR+): P < 0.001 (***), n = 9 mice; BBS (NIR+) vs levodopa: P = 0.031 (*), 9 mice in the BBS (NIR+) group and 6 mice in the levodopa group; BBS (NIR+) vs Pre-lesion: P = 0.128 (n. s.), 9 mice in the BBS (NIR+) group and 6 mice in the pre-lesion group. (**j, k**) Print area (**j**) and stride length (**k**) for each foot of the PD mice implanted with the BBS in the CatWalk gait analysis before (NIR-) and after (NIR+) the NIR irradiation. Print area: BBS (NIR+) vs BBS (NIR-): RF: P = 0.008 (**), RH: P = 0.013 (*), LF: P = 0.028 (*), LH: P = 0.028 (*); n = 8 mice. Stride length: BBS (NIR+) vs BBS (NIR-): RF: P = 0.016 (*), RH: P = 0.010 (*), LF: P < 0.001 (***), LH: P = 0.036 (*); n = 8 mice. RF: right front paw, RH: right, hind paw, LF: left front paw, LH: left hind paw. (**l**) Statistics of regularity index in the CatWalk gait analysis for the health mice (Pre-lesion), the PD mice implanted with the cell-laden devices (Cell-laden), the cell-free devices (BS), or the BBS before (NIR-) and after (NIR+) the NIR irradiation, and the PD mice treated with levodopa. BBS (NIR+) vs BBS (NIR-): P < 0.001 (***), n = 8 mice; BBS (NIR+) vs BS: P < 0.001 (***), n = 8 mice; BBS (NIR+) vs Cell-laden: P < 0.001 (***), n = 8 mice; BBS (NIR+) vs levodopa: P = 0.041 (*), 8 mice in the BBS (NIR+) group and 6 mice in the levodopa group; BBS (NIR+) vs Pre-lesion: P = 0.965 (n. s.), 8 mice in the BBS (NIR+) group and 6 mice in the pre-lesion group. (**m**) Representative MRI images of mouse brain with implanted BBS for 60 and 120 days. The implanted BBS is highlighted by white arrows. Scale bar: 2 mm. (**n**) Representative immunofluorescence images of brain tissues with the implantation of the BBS at post-surgery 120 days with stained GFAP (red, for labelling activated astrocytes related to gliosis), NeuN (green, for labeling neurons), and nucleus (blue). Scale bar: 100 μm. (**o**) Statistics of neuronal density and fluorescence intensity of GFAP ipsilateral to the implantation of the BBS over different time periods. neuronal density: Day 30 vs Day 120: P = 0.951 (n. s.); GFAP fluorescence intensity: Day 30 vs Day 120: P = 0.012 (*). n = 5 separate experiments. Statistical analyses into the data shown in **d, j, k**, and **o** were performed using one-way ANOVA with two-tailed Student’s *t*-test. Statistical analyses into the data shown in **f, g, i**, and **l** were performed using one-way ANOVA followed by a post hoc with Bonferroni correction for multiple corrections.

Following BBS implantation and NIR irradiation, a significant increase in c-Fos-positive neurons, a marker of neuronal activation, is observed in the midbrain compared to the controls, including pre-irradiation and cell-free brain stimulators groups (Fig. 4c, d). These findings confirm that the BBS can functionally integrate with natural neural circuits and evoke physiological neural signalling in vivo. Additionally, the numbers of TH-positive neurons are markedly higher in the ipsilateral midbrain relative to the contralateral, non-implanted site (Fig. 4c). Remarkably, the BBS induces an approximately 9-fold increase in TH intensity compared to cell-free brain stimulators (Fig. 4d, Fig. S18a, b), suggesting potential restoration of endogenous neural function and facilitation of dopamine neurotransmission efficiency^54^.

We next evaluate the restoration of locomotor function in PD mice by conducting behavior tests on days 14 and 18 post-implantation. After NIR irradiation, PD mice implanted with the BBS exhibit significant improvements in general locomotor activity, as evidenced by increased distance traveled and higher movement speed in open-field tests compared to pre-treatment conditions (Fig. 4e-g and Fig. S18c-e). In rotarod tests, BBS-treated mice demonstrate enhanced motor skills, remaining on the rod for longer durations after NIR irradiation (Fig. 4h, i and Fig. S18f). Furthermore, CatWalk tests reveal improved gait- and balance-related dynamic coordination, as indicated by reduced print area, increased stride length, and a high average regularity index (∼96%), comparable to pre-lesion levels (Fig. 4j-l and Fig. S18g, h). In contrast, no significant improvements in locomotor function are observed in mice implanted with cell-laden control devices or cell-free brain stimulators under identical NIR conditions (Fig. S19, Fig. S20). Across open-field, rotarod, and Catwalk tests, BBS treatment outperform the “gold-standard” levodopa therapy (Fig. 4f, g, i, l, and Fig. S21), providing sustained, controllable dopamine for over 336 hours and overcoming the limited therapy window of levodopa (∼6 hours)^55^.

To evaluate long-term efficacy, the BBS is implanted in PD mice for 120 days (Fig. S22a). The BBS maintains stable micropyramid structures, consistent electric field generation, and preserves ∼18% viable cells (Fig. S22b-g), substantially higher than that observed with direct cell transplantation (∼6% viability after 90 days)^56^. Behavioral assessments confirm persistent therapeutic benefits in locomotor activity, motor skill, and gait coordination under NIR irradiation, whereas removal of the BBS results in the recurrence of Parkinsonian symptoms (Fig. S22j-n). These results demonstrate the long-term in vivo effectiveness of the BBS and its therapeutic potential for PD.

Finally, we assess the biosafety of the BBS. Though slight neuroinflammatory cell infiltration at acute time points (21 and 30 days post-implantation), no pathological cavities formed at the implantation sites, and scar tissue gradually resolves over a prolonged implantation period of 120 days (Fig. 4m, Fig. S23, and Fig. S24a). The foreign body response is negligible, as evidenced by declining glial cell activation and stable neuronal densities adjacent to the BBS (Fig. 4n, o). After 120 days, the densities of glial cells and neurons surrounding the BBS are comparable to these in contralateral, non-implanted regions (Fig. S24c, d). In contrast, conventional deep brain stimulators induce pronounced gliosis and significant neuronal loss at implantation sites, leading to side effects^57^. Histological analysis of major organs (brain, kidney, spleen, liver, heart, and lung) reveals no pathological changes following long-term BBS implantation (Fig. S24e). Notably, BBS implantation has no detectable impact on locomotion function in healthy mice, indicating an absence of dyskinesia or other side effects (Fig. S25). Collectively, compared to existing therapeutic approaches (*e*.*g*., levodopa, brain stimulation, and cell therapy), the BBS that integrates the merits of both bioelectronics and medicine offers superior, synergistic efficacy in reversing bradykinesia, as well as motor and gait deficits in PD, while eliminating long-term biosafety concerns and complications commonly associated with these therapies (Fig. S24f-I, Fig. S26, and Table S2)^5, 55, 58^.

## Conclusions

In summary, we introduce a living bioelectronic medicine-BBS that harnesses the advantages of both bioelectronics and medicine by integrating ferroelectric bioelectronics, dopaminergic cells, and gelatin hydrogel, achieving neurotransmitter-evoked neural signaling. Unlike conventional implanted brain stimulators, this BBS are capable of programmed secretion of physiological-level dopamine (∼7 ng·mL^-1^, equivalent to the dose in normal rat’s striatum), imparting the unique capability of specific activation of dopaminergic signalling both in vitro and in vivo. Following long-term implantation in PD mice, this lifelike BBS not only effectively restores motor function but also exhibits no detectable side effects and negligible foreign-body responses, moving an important initial step toward practical applications. Eliminating biosafety and therapeutic efficacy concerns associated with conventional strategies, this living bioelectronic medicine will provide a clinically viable route for neuromodulation, holds great promise for next-generation bioelectronics, medicine, and brain-machine interfaces^29-32, 59, 60^.

## Acknowledgments

The authors acknowledge the financial support provided by the National Natural Science Foundation of China (52022102, 52173148, and 52261160380), the Shenzhen Medical Research Fund (B2302045), National Key R&D Program of China (2017YFA0701303), the Youth Innovation Promotion Association of CAS (Y2023100, 2022368), Guangdong Regional Joint Fund-Key Project (2021B1515120076), the Fundamental Research Program of Shenzhen (RCJC20221008092729033, JCYJ20220818101800001). The authors thank Professor Yu-Tian Wang and Doctor Anping Chai for their great help in whole-cell patch-clamp recording. The authors also thank Professor Yi Lu and Ms. Yiyong Wu for their great help in amperometric dopamine recording. Professor Hao Qian from University of Electronic Science and Technology of China is also acknowledged for his constructive suggestions and discussion.

## Author contributions

X. D conceived the idea and supervised the research. Q. Z., Q. M., and M. P. conducted the experiments with the assistance of Z. L., Z. L., X. J. and H. Z. Q. Z. and X. D analyzed the results and wrote the manuscript. All authors contributed to the discussion and interpretation of the results. Q. Z., Q. M. and M. P. contributed equally to this work.

## Data availability

All data presented in this work are available from the corresponding authors upon reasonable request.

## Supplementary Materials

Materials and Methods

Supplementary Figures S1 to S26

Supplementary Tables S1 to S2

Supplementary Videos 1 to 18

